# Ethanol reduces grapevine water consumption by limiting transpiration

**DOI:** 10.1101/2024.08.31.610611

**Authors:** Neila Ait Kaci, Alice Diot, Beatrice Quinquiry, Olivier Yobregat, Anne Pellegrino, Pierre Maury, Christian Chervin

## Abstract

Studies suggest that ethanol (EtOH), triggers plant adaptation to various stresses at low concentrations (10 µM to 10 mM). This study investigates whether EtOH induces drought acclimation in grapevine, as demonstrated previously in *Arabidopsis*, rice, and wheat. Preliminary results with bare root Gamay cuttings showed that those pre-treated with 10 µM EtOH aqueous solutions lost fewer leaves when deprived of water compared to controls. Subsequently, we ran a potted-cutting experiment with progressive soil water deficit. Plants pre-treated with 250 mM EtOH solutions exhibited slower depletion of the fraction of transpirable soil water (FTSW), compared to controls. While 250 mM EtOH decreased transpiration in early days, these EtOH pre-treated plants maintained higher leaf transpiration than controls after 10 days of soil water depletion. The transpiration response to FTSW was affected by EtOH application. EtOH pre-treatments limited plant leaf expansion without increasing leaf senescence. Interestingly, plants primed with EtOH followed typical hormesis curves. These results suggest that EtOH improves grapevine acclimation to drought, leading to potential water-savings in wine growing regions prone to high water shortages, linked to climate change. These should encourage further testing under various vineyard conditions.

## Introduction

Many recent studies show that ethanol at physiological concentrations can have positive roles in plant development (Diot *et al*., 2024a). In particular, the work of Bashir *et al*. (2022) demonstrated that ethanol could induce drought acclimation in several plant species, including Arabidopsis, rice, and wheat. This induced acclimation is particularly interesting in the context of climate change, which is expected to increase water stress in certain geographical areas, such as the Mediterranean region (Cramer *et al*., 2018; IPCC, 2021). Given that viticulture is particularly developed in this region (OIV, 2022), we sought to test the effect of ethanol on grapevine under controlled conditions of water stress, to verify if the observations of Bashir *et al*. (2022) are applicable to this species.

Initially, experiments were conducted on grapevine cuttings with bare roots in water, with the main observation being the percentage of dried leaves one week after rehydration. Then, trials were carried out on potted grapevine cuttings, monitoring the weight of the pots to estimate plant transpiration (Rodrigues *et al*., 2019). This gravimetric method also enabled the estimation of the fraction of transpirable soil water (FTSW), a variable widely used to quantify soil water deficit imposed on the plant and its impact on the plant main functions (Sinclair, 2005). Yet, leaf expansion exhibits the highest sensitivity, followed by plant transpiration, and leaf senescence; this sensitivity to soil drying is assessed by the FTSW threshold (Kang *et al*., 2024). Our innovative results about ethanol altering grapevine transpiration and functional leaf area lead us to examine our RNA-seq data obtained in a parallel study regarding grapevine resistance to heat stress (Diot *et al*., 2024b). We hope to quickly stimulate further research to find the appropriate conditions of EtOH application for the various varieties and geographical situations encountered in vineyards affected by water stress episodes.

## Material and Methods

### Plant material

Grapevine cuttings were made from one-year-old dormant Gamay shoots (*Vitis vinifera*), clone 787, from a collection of genetic material whose plant material is tested to be safe for the main vine viruses (IFV Sud-ouest, Peyrole, France). The cuttings were prepared as described by Mullins (1966), with the following modifications. Single-node cuttings, 5-cm long with two cutter scars at the bottom to facilitate rooting, were disinfected with active chlorine (1.5% v/v) for one minute and rinsed three times with running water. Then, 12 cuttings were placed in 250 ml plastic crates filled with 150 ml water and covered with aluminium foil to hold the cuttings up, the lower end being immersed in water, the bud ending up upwards (Fig. 1A). The cuttings were left for one month in a growing chamber at 25°C/20°C 16h/8h (day/night) with light set to 250 µmol.m^−2^.s^−1^. The water level was maintained to the initial filling mark.

**Fig. 1.**
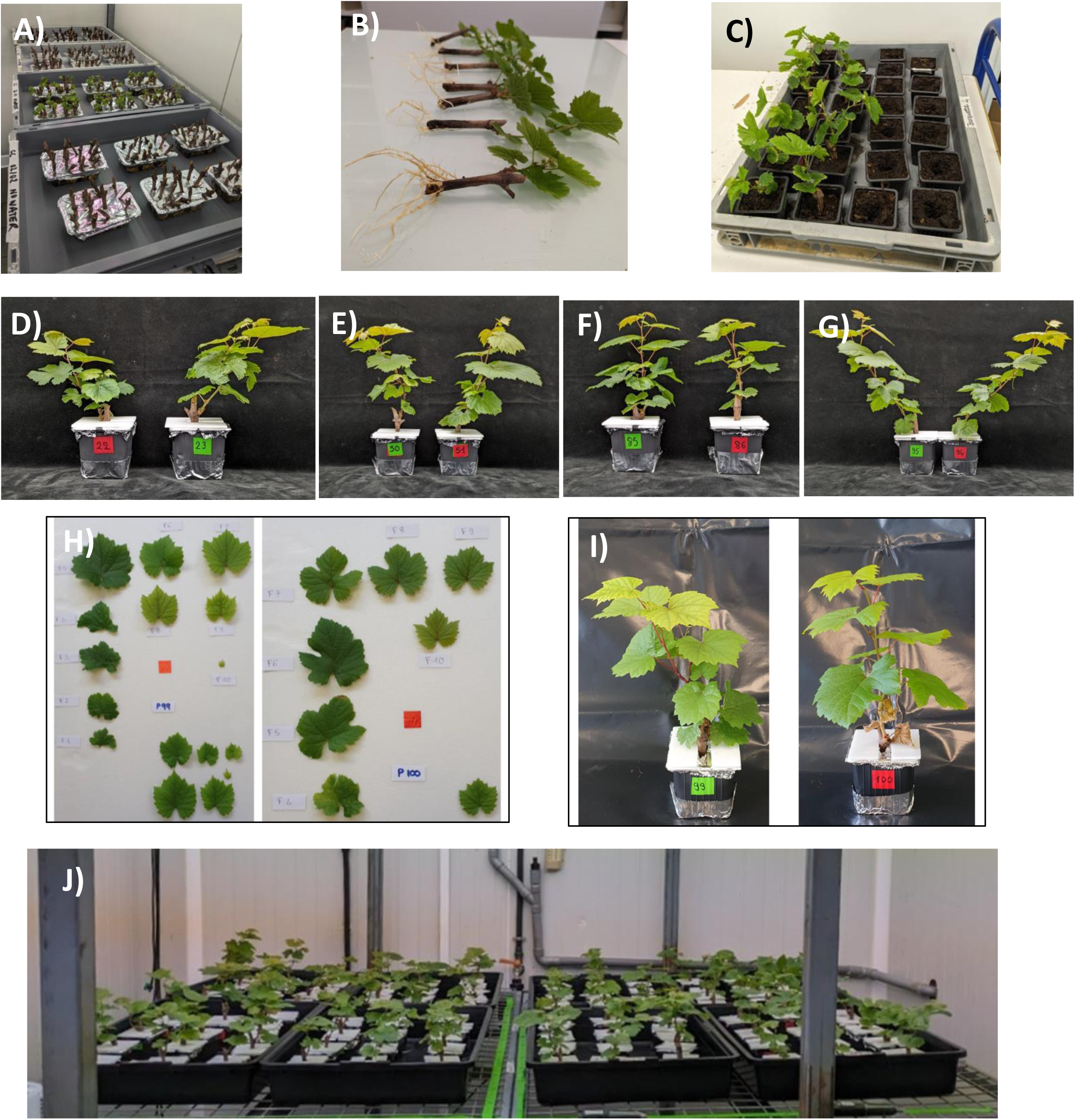
Various stages of the culture of Gamay cuttings: A) bare root cuttings in water; B-C) repotting after one month; D-G) variability of cuttings; H) measurement of leaf surface with Easy Leaf Area; I) example of dry leaves on the bottom at the end of the experiment; J) set-up in the growth chamber.

Two experiments (Exp. 1 & Exp. 2) were conducted. Exp. 1 was based on bare root cuttings with 4 biological replicates of 6 plastic crates, one per EtOH concentration with 12 cuttings per crates, thus a total of 288 cuttings. Exp.2 was performed with potted cuttings with 8 biological replicates of 6 potted cuttings, one potted cutting per EtOH concentration, two water treatments, thus a total of 96 cuttings in pots.

### Ethanol treatments applied to bare-root cuttings

In Exp. 1, the water was removed and 150 ml of freshly prepared ethanol solutions at various concentrations (0, 0.01, 0.1, 1, 10 and 100 mM) were poured into the crates and left for 24 h to allow ethanol priming. Then, all solutions were removed and the cuttings were left without water for 38h (duration determined by preliminary trials). After this drying period, 150 ml of water were poured into each crate, and the cuttings were left to recover in the growing chamber. The percentage of dry leaves was assessed one week later. An example of what we considered as dry leaves is shown in Fig.1I.

### Ethanol treatments applied to potted cuttings

In Exp. 2, the one-month cuttings (Fig. 1B, C) were transferred into 105 ml nursery plastic pots, filled with P.A.M.2 Proveen substrate (Proveen, Netherlands), and water saturated. Plants were left to settle down and develop roots for one week in a growing chamber 25°C/20°C (day/night) with a 16 h photoperiod providing 250 µmol.m^−2^.s^−1^ photosynthetically active radiation at the top of the plants. The plants were maintained at maximum Fraction of Transpirable Soil Water (FTSW = 1) over the week. The pots were covered with aluminium foil on the top, and an additional 3 mm polystyrene sheet (Depron, Germany) was added onto the top to limit soil evaporation (Fig. 1D, E, F, G, J). Evaporation did not exceed 10% of the FTSW after 30 days (Supplementary Fig. S1). Then, the day prior to the beginning of the water deficit, 52.5 ml of ethanol solutions at various concentrations were poured onto the pots, after removing temporarily the bottom aluminium foil to allow run-off of the solution into a saucer. The ethanol concentrations tested in this study were 0, 0.4, 2, 10, 50 and 250 mM. These concentrations correspond to ethanol/water ratios of 0, 0.002, 0.012, 0.06, 0.29, and 1.46% (v/v), respectively. The range was chosen with 5-fold increments to minimise errors from soil variability. Pairs of plants one well-watered (WW) and one water-stressed (WS) were established for all EtOH doses. A total of eight blocks were set-up, in each block we added one pot without any plant, to measure the evaporation of the pot.

### Soil water status measurements

The day following the ethanol treatment (Exp. 2), the watering of Water Stress “WS” pots was stopped, while the Well Watered “WW” pots were maintained at FTSW = 1. FTSW was calculated from pot weights, as described previously (Lebon *et al*., 2006). The daily FTSW was calculated from pot weight on d day (pot weight d), the pot weight at full soil water capacity (pot weight at fc), and the pot weight when transpiration of the non-irrigated pot was less than 10% of WW controls of the same EtOH treatment (pot weight 10%). Prior to water deficit treatment, pots were irrigated to saturation, and after the excess water had drained, the pots were weighed to estimate the pot mass at full soil water capacity. Each pot was weighed daily, twice for the WW, before and after re-watering. FTSW_d_ was then determined as follow (eq.1):

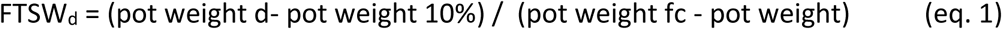

### Grapevine transpiration and leaf area measurements

Water loss for individual plants (T) over 24h (Exp. 2) was calculated from the pot weights difference, considering pot weight after irrigation (d-1 day) and before irrigation (d day) for WW. Then, transpiration rate per leaf area (Tr) was determined by dividing the daily pot weights difference by the total leaf area values measured as close as possible to the transpiration date (eq.2).

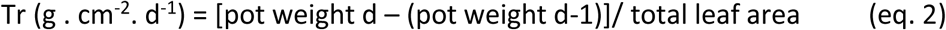

The leaf area of all plants was estimated using a regression between the longest leaf vein and the leaf area of 230 leaves (Supplementary Fig. S2). The leaf area was determined with the “easy leaf area” application (https://www.quantitative-plant.org/software/easy-leaf-area). For this purpose, leaf main vein lengths were measured twice, once around day 3 and once around day 8. The calculation of plant transpiration of days 2, 3, 4 and 5 was performed using the first set of leaf area estimates, and the transpiration of days 6, 7, 8, 9 and 10 was calculated using the second set of leaf estimates.

### Normalised transpiration rate (NTR), and NTR response to soil water status

Daily plant transpiration values (T in g . pot^−1^. d^−1^) from day 1 to around day 13 (except for 250 mM up to day 24) were normalised as described by King and Purcell (2017), according to equation 3. Briefly, a correction factor (CF) was calculated to adjust for differences in transpiration among plants within an EtOH treatment under WW conditions. The correction factor was calculated while all plants were WW (before Day 0) by dividing the water loss of individual plants by the average water loss of the WW plants at the same EtOH dose. A second correction was made to account for differences in transpiration that occurred among days after initiation of water deficit treatment. Water loss for individual plants was divided by the average water loss for WW plants of the same EtOH treatment (T_WW average). NTR for each plant for a given weighing interval was:

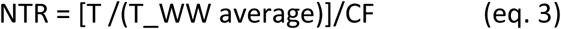

Then the response of NTR at the whole plant level to FTSW was fitted according to equation 4 (Muchow and Sinclair, 1991), in which “*a*” is the model parameter describing the response of NTR to FTSW, “*a*” characterise the curvature of the fit. As the “*a*” parameter increases, the sensitivity of the NTR strengthens at a given same FTSW value. The FTSW threshold (FTSWt) represents the FTSW at which NTR (modelled by equation 4) starts to decrease (5% of reduction compared to maximum NTR, as proposed for nonlinear models by Sadras and Milroy, 1996). A higher FTSWt indicates a higher sensitivity to water deficit (Andrianasolo *et al*., 2016).

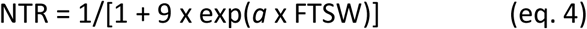

### Re-watering of WS plants and experiment ending

After preliminary tests, the re-watering time for the WS plants was when at least two top leaves in the canopy began to wilt. The plants were re-watered with 30 ml the first day, then the amount of water was calculated daily to reach FTSW = 0.95. The plants were harvested one week after re-watering. The WW plants and their corresponding WS plants (being part of the same pair, as defined above) were harvested the same day. The harvest period was spread over a week from day 28 to day 34 after the beginning of Exp. 2. On the harvest day, all leaves and roots were collected and the leaf area was estimated directly with Easy Leaf Area application. The percentage of dried leaves was calculated over the seven oldest leaves, as at day 0 there was a minimum of seven leaves per plant (example of dry leaves in Fig. 1I). Finally, the roots were dried at 60°C for at least 72h, and then weighed.

### RNA-seq analyses on grapevine cell cultures

RNA-seq analyses have been performed on *Vitis vinifera* cv. Gamay Fréaux cell cultures. The complete material and method is described in Diot *et al*. (2024b). However, the following modifications in the formal analysis have been done: the relative estimated transcript counts have been performed only with the two conditions used in this study (comparison between 6h_0mM and 6h_1mM) and for this analysis, the thresholds for Differentially Expressed Genes were set at |Log2 Fold Change| > 0.5, and padj < 0.1. The raw data discussed in this publication have been deposited in NCBI’s Gene Expression Omnibus (Edgar *et al*., 2002) and are accessible through GEO Series accession number GSE275842 (https://www.ncbi.nlm.nih.gov/geo/query/acc.cgi?acc=GSE275842).

### Statistical treatments

All fitting and statistical analyses were performed with Rstudio version 4.2.1, using the packages ‘car’ and ‘rstatix’. Data normality and homoscedasticity hypotheses were first assessed using Shapiro-Wilk tests and Brown-Forsythe tests, respectively. According to the cases, pairwise comparisons were performed with Student’s tests, Welch’s tests or Wilcoxon’s tests. Fits of NTR to FTSW were performed using the non-linear least squares (NLS) regression model. The goodness of fit was assessed by calculating the Rsquare (R^2^), root mean square error (RMSE), and mean squared error (MSE).

## Results

### Ethanol priming reduced the percentage of dried leaves in bare root cuttings subjected to a temporary water depletion

A preliminary experiment (Exp.1) was conducted with the bare root cuttings, by priming them with EtOH at various concentrations, before removal of the aqueous solutions for 38h, then re-filling the crates with water. After an additional week, we observed the amount of dried leaves (Fig. 2). Although not significant, there was a clear trend of reduction of the dried leaves for plants primed with EtOH compared to controls. Indeed, the priming with EtOH reduced the percentage of dried leaves from 35.5% (controls at 0 mM) to 17.7% (EtOH at 0.01 mM). Instead of running more bare root trials to reach significance levels, with conditions which are not natural, we decided to run a larger trial, with cuttings in light potting soil (Exp. 2), and to follow a larger set of parameters. At least, these conditions can be met in some nurseries.

**Fig. 2:**
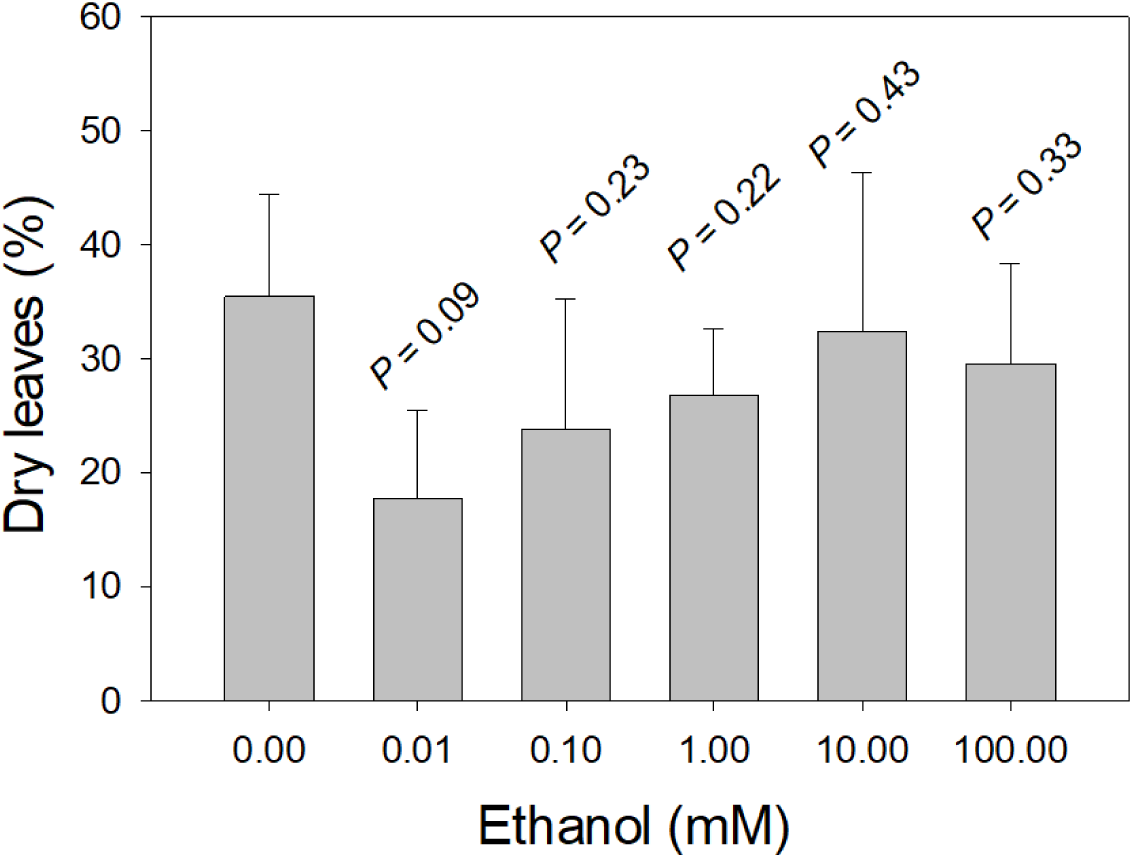
Effects of ethanol (EtOH) priming on Gamay one-node cuttings with bare roots (preliminary study). One-month-old cuttings were primed with EtOH at various concentrations for 24h, then left without water for 38h, then assessed for dry leaves one week after rewatering; 4 replicates of 12 cuttings per concentration. Pairwise comparisons with one-tail t-tests against controls (0).

### Ethanol maintained higher Fractions of Transpirable Soil Water than controls both for Well Watered and Water Stress pots

The changes in Fraction of Transpirable Soil Water (FTSW) over a three-week period in potted single-node cuttings (Exp. 2) were plotted for WW pots (Fig. 3A) and WS pots (Fig. 3B). FTSW of WW pots were brought up daily to 0.95. The results show the FTSW just before re-watering the pots. Most WW pots remained above 0.7, regardless of the EtOH concentration used for priming. Occasionally, 250 mM plants showed significantly higher values than controls (Fig. 3A and Supplementary Table S1). On the other hand, the FTSW of WS pots gradually decreased after their last irrigation on day 0. In plants primed with 250 mM EtOH, the FTSW decreased significantly more slowly than in control conditions (Fig. 3B and Supplementary Table S1). Finally, given that there was a negligible decrease in FTSW in pots without plants due to soil evaporation (Supplementary Fig. S1), we assumed that changes in FTSW were primarily due to plant transpiration and tested this hypothesis in a series of subsequent analyses.

**Fig. 3.**
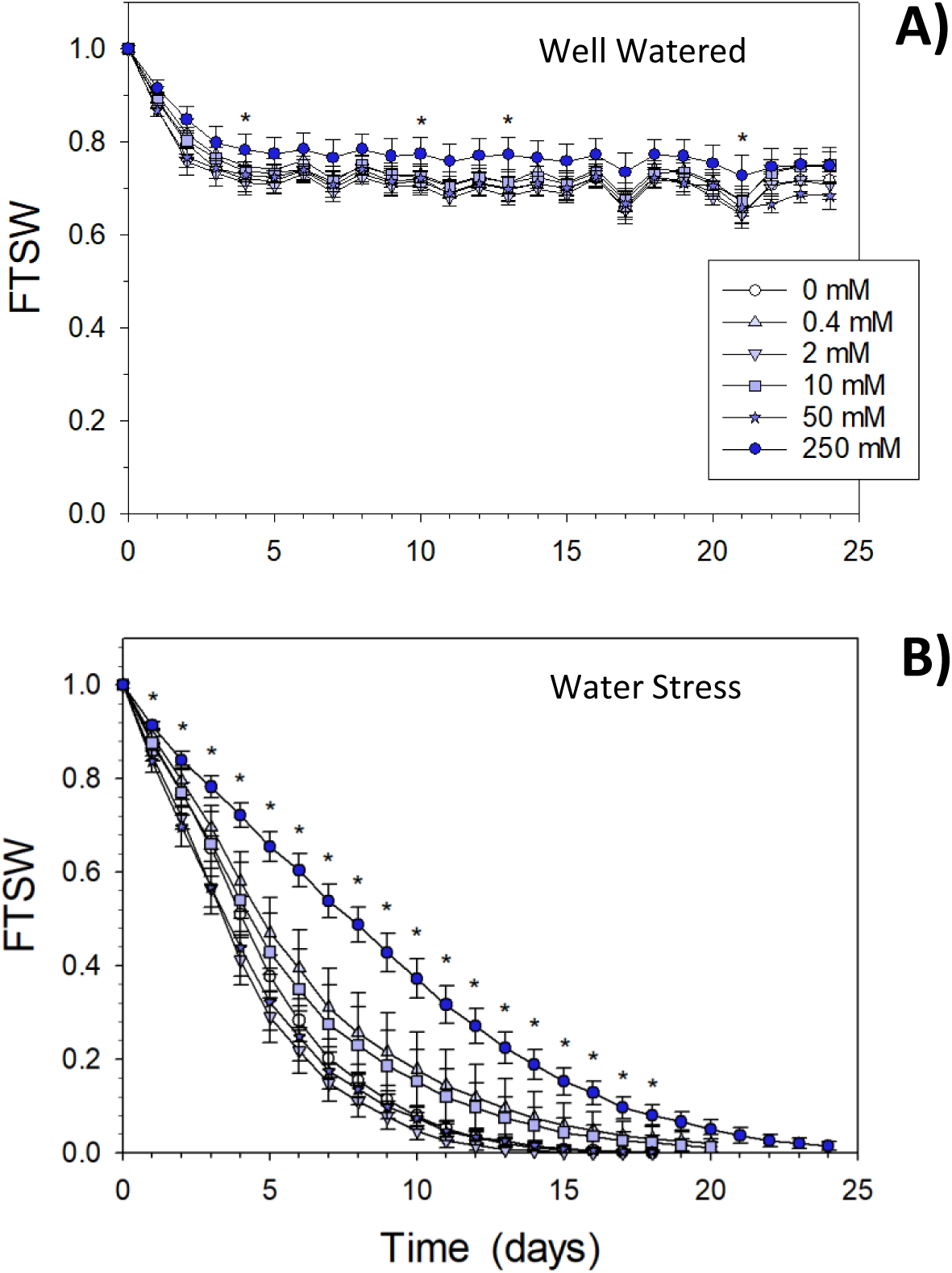
Effects of ethanol (EtOH) priming on FTSW as a function of time after « soil water deficit » initiation in Gamay one-node cuttings growing on light potting soil. **A)** well-watered plants, WW, **B)** water stress plants, WS, irrigation stopped at day 1. n = 8, error bars show S.E. * stands for significant differences at 0.05 between 0 and 250 mM (details in Supp. Table S1).

### Ethanol slowed down the grapevine transpiration

Figure 4 shows the changes in transpiration per unit of leaf area from day 3 to day 10, a period with the biggest FTSW differences between the various treatments (Fig. 3). The transpiration of WS plants started to be significantly lower than WW plants from day 6, as shown with asterisks (Fig. 4; Supplementary Tables S2 and S3). It is noticeable that 250 mM EtOH reduced significantly plant transpiration, in comparison to controls, from day 3 to day 5 in WS plants, and from day 3 to day 10 in WW plants. We also observed that, compared to the controls, the transpiration of WS plants treated with EtOH showed opposite trends at the beginning and at the end of the experiment. For example, cuttings treated with 250 mM of EtOH had a lower rate of transpiration than controls at days 3, 4 and 5, but displayed higher transpiration than controls at days 8, 9 and 10 (Fig. 4 and Supp. Table S2). Interestingly, when pooling the transpiration results from day 2 to day 6, a significant transpiration decrease in the plants treated with 0.4 mM EtOH compared to controls was also noticed (data not shown). Finally, we observed a bell-shaped distribution in the transpiration rates according to increasing EtOH doses (not taking the controls with no EtOH priming into account), particularly noticeable in WW and WS at day 10 (Supplementary Fig. S3A and S3B), but also at days 7, 8 and 9. This is classical of a hormesis effect, which will be discussed later.

**Fig. 4.**
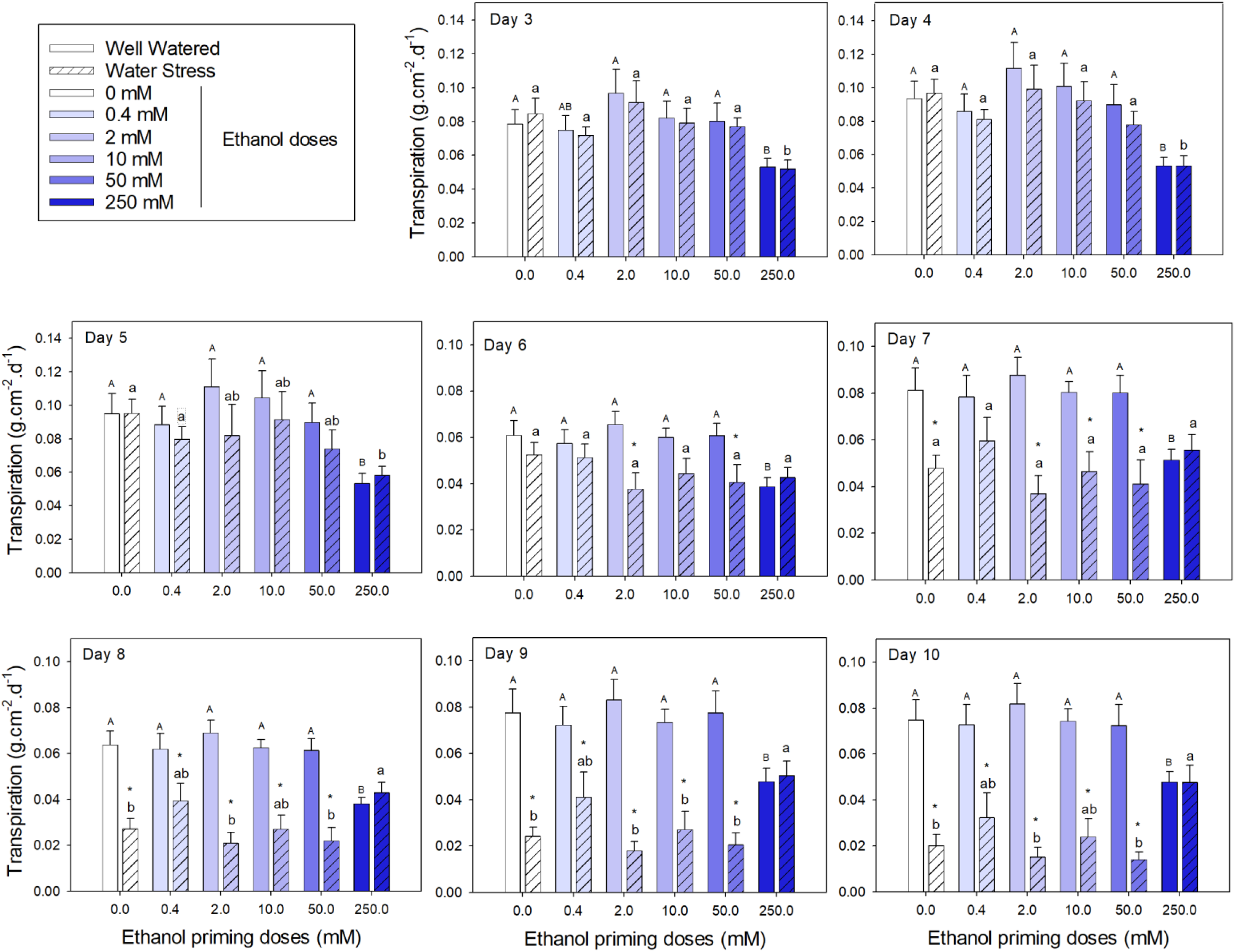
Effects of ethanol (EtOH) priming on transpiration (Tr) of Gamay one-node cuttings in pots, from day 3 to day 10 after the initiation of soil water deficit n = 8, error bars show S.E.; all statistical details in Supp. Tables S2 and S3; multiple comparisons: capital letters to compare within WW; small caps to compare within WS; * stands for significant differences at 0.05 between WW and WS within the same EtOH dose.

### Ethanol altered the plant transpiration response to FTSW

The normalised transpiration rate (NTR) response to FTSW is shown in Figure 5. NTR started at 1 (to the right of the graph) when the plants transpired at the same rate as the WW plants within the same EtOH treatment, then dropped to lower values (to the left of graph) when FTSW became a limiting factor for transpiration. According to the values of R^2^, RMSE, MSE (Table 1), the performance of non-linear fitting was satisfactory. The FTSW threshold (FTSWt) from which NTR dropped below 0.95 was determined. We observed that priming plants with 250 mM led the plants to switch the FTSWt to higher values (FTSWt = 0.67) than plants primed with lower doses (FTSWt < 0.50) (detailed values and stats are given in Table 1 with *p*-values detailed in Supplementary Table S4). Regarding the “*a*” parameter, the curvature coefficient, pairwise comparisons showed significant differences between 0.4 mM and 50 mM, compared to 250 mM. Thus, priming plants with 250 mM EtOH showed earlier regulation of transpiration in response to a progressive soil drying.

**Fig. 5.**
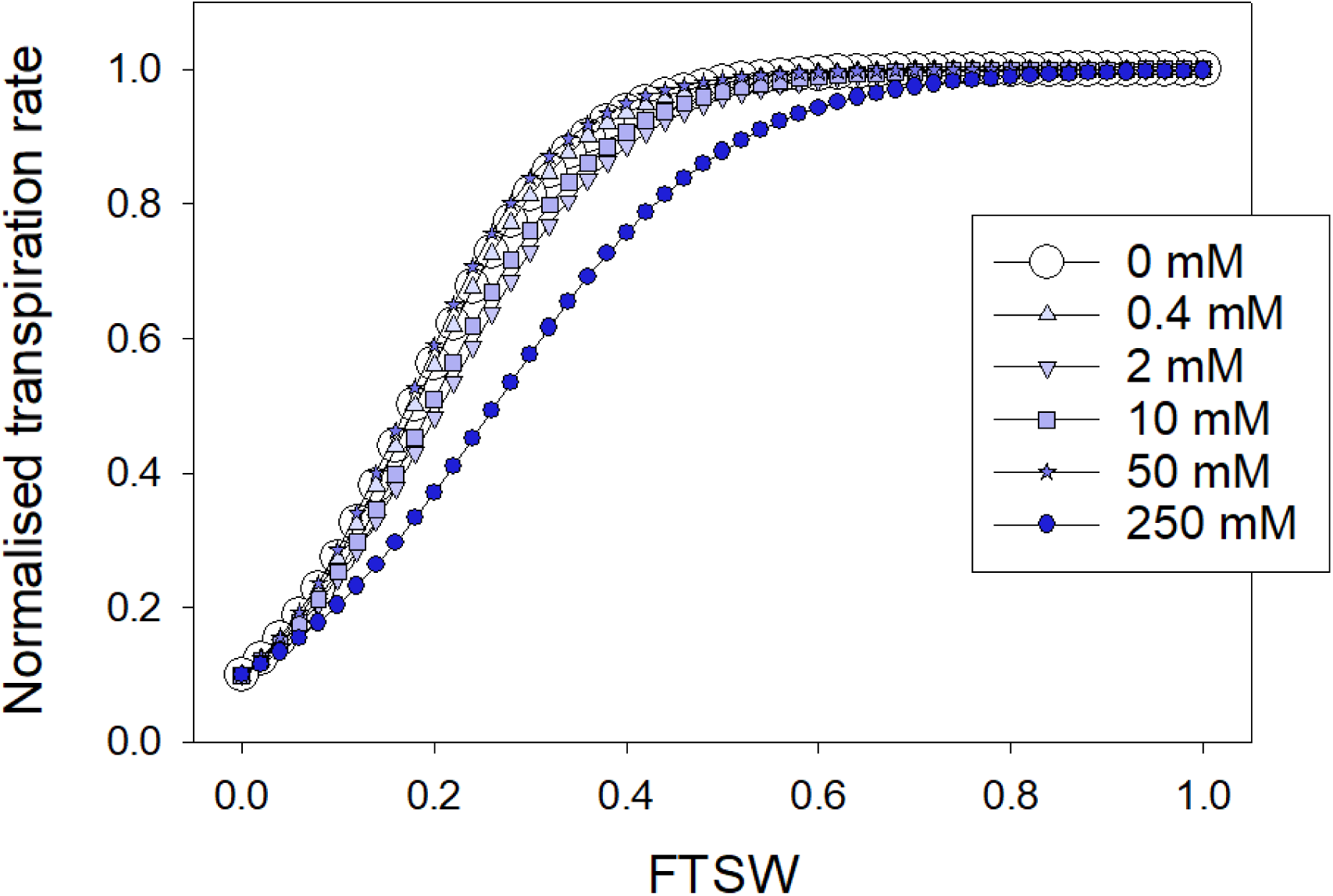
Fitted response of normalised transpiration rate (NTR) at the whole plant level to the fraction of transpirable soil water (FTSW) for six ethanol priming doses 0, 0.4, 2, 10, 50 and 250 mM. The curve fitting and calculations have been performed with the R software as described in Material and Methods (see fitting quality parameters in Table 1).

**Table 1.** Parameters describing the normalised transpiration rate (NTR) at the whole plant level to the fraction of transpirable soil water (FTSW) variations for six ethanol (EtOH) priming doses. All fitting and calculations details are given in Material and Methods. The “a” parameter represents the curvature of the fit; FTSWt stands for FTSW threshold at which NTR maximum was reduced by 5%. R² is the goodness of fit. RMSE stands for root mean square error. MSE stands for mean square error. Significant differences were determined at *p* < 0.05 using pairwise comparison methods (see Material and Methods and detailed *p*-values in Supp. Table S4).

| EtOH dose (mM) | "a" | Signif. | FTSWt | Signif. | $R^2$ | RMSE | MSE |
| --- | --- | --- | --- | --- | --- | --- | --- |
| 0 | -12.02 | a | 0.44 | a | 0.95 | 0.095 | 0.009 |
| 0.4 | -12.50 | a | 0.43 | a | 0.91 | 0.129 | 0.017 |
| 2 | -10.96 | ab | 0.50 | ab | 0.94 | 0.105 | 0.011 |
| 10 | -11.38 | ab | 0.50 | ab | 0.87 | 0.141 | 0.020 |
| 50 | -12.81 | a | 0.41 | ab | 0.92 | 0.168 | 0.028 |
| 250 | -8.60 | b | 0.67 | b | 0.87 | 0.133 | 0.078 |

### Soil water deficit and ethanol differentially impacted the leaf development and senescence

As shown in Figure 6 the soil water deficit limited the leaf expansion over the 4-week trial. At the end of the experiment, the average leaf area of the WW control plants was just above 200 cm² (Fig. 6), when the one of the WS control plants was just above 100 cm² (Fig. 6). The EOH priming also reduced the leaf expansion, this was particularly noticeable on the plants primed with 250 mM EtOH which were harbouring an average leaf area twice smaller than the controls (0 mM EtOH) in WW plants and in WS plants (Fig. 6). The results of Figure 7 showed that, at harvest, WW plants harboured around 10% of dried leaves, while WS plants harboured around 80% of dried leaves (Fig. 7). In none of these two series, WW and WS, the ethanol changed significantly the % of dried leaves.

**Fig. 6.**
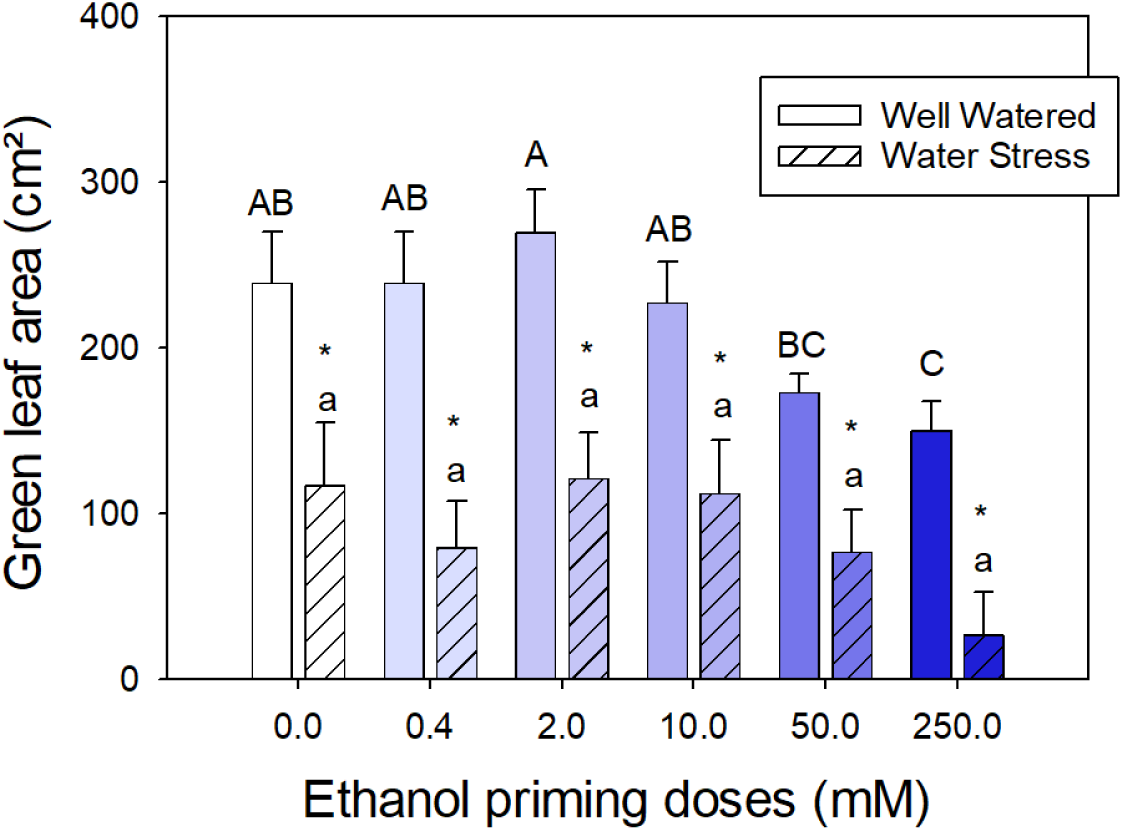
Impact of ethanol (EtOH) priming on leaf area at harvest, n = 7 plants, error bars show SE. Multiple comparisons details in Supp. Table S5. Capital letters to compare within WW, small caps to compare within WS, and * shows *p* < 0.05 when comparing water deficit effect within a similar EtOH dose.

**Fig. 7:**
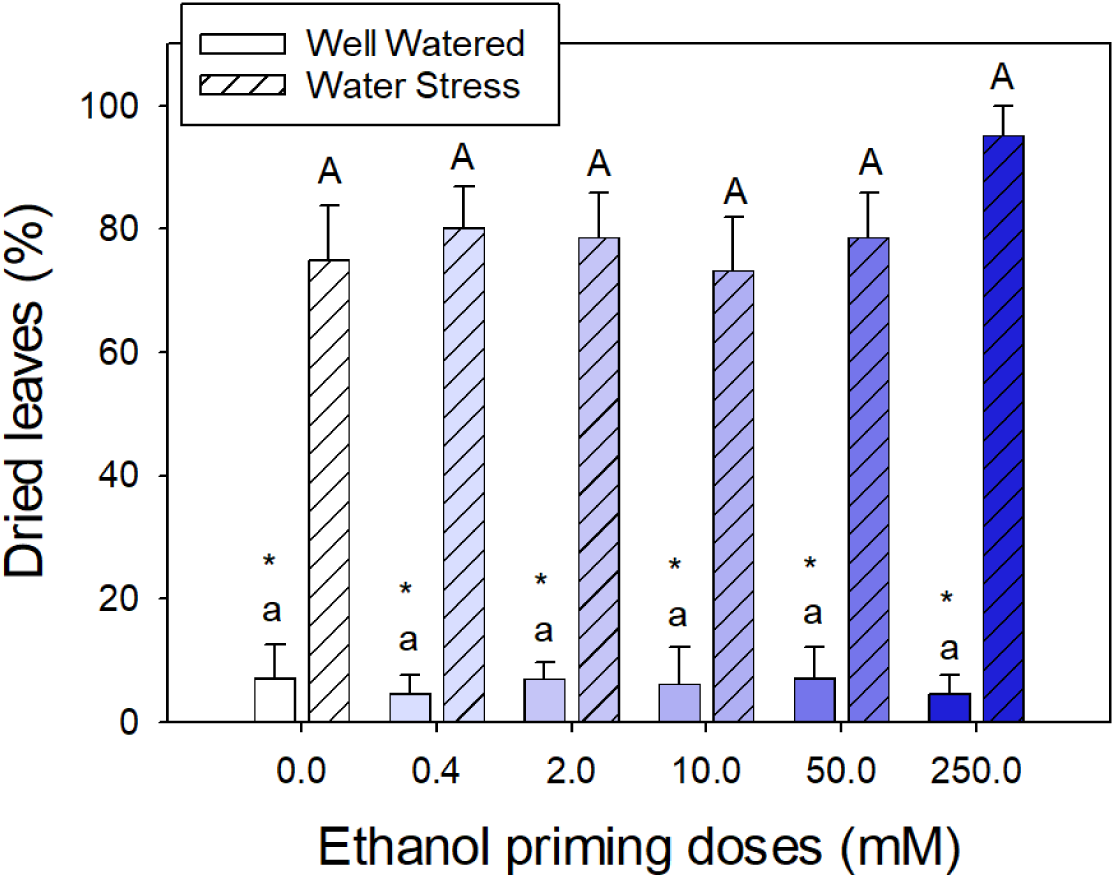
Impact of ethanol (EtOH) priming on the % of dry leaves at harvest, n = 7 plants, error bars show SE. Multiple comparisons details in Supp. Table S6, * shows *p* < 0.05 in all comparisons of water deficit effect within each EtOH dose.

## Discussion

### Ethanol priming and grapevine bare root cuttings

Water is becoming a scarce resource and the risk of excessive drought linked to climate change is a serious concern for vineyard management (van Leeuwen *et al*., 2024). Ethanol priming at very low doses (0.4 to 250 mM) may provide a better acclimation of grapevine to drought, notably by limiting transpiration while maintaining functional leaf tissues. These ethanol doses were close to physiological values measured in Gamay cells, 100 µM to 2 mM (Diot *et al*., 2024b). In a preliminary experiment with bare root cuttings, exposed to air for more than 30 hours, we observed a reduction in dry leaves in the cuttings pre-treated with 10 µM EtOH, one week after re-watering, compared to non-treated plants. Such a concentration corresponds to 0.0006% of ethanol in water, which is extremely low. However, it is hypothesised to be in the range of detectable level of EtOH for organisms, possibly used as a signal to adapt to stressful environments, by eliciting physiological changes (Diot *et al*. 2024a).

### Ethanol reduces water use by potted cuttings

These preliminary results on bare root cuttings prompted us to carry out more physiological tests with plants in soil, in order to apply water deficit kinetics on root systems closer to vineyard conditions. The FTSW dropped from 1 to less than 0.1 after 15 days, on average, in most water-deprived plants (Fig. 3B), which is consistent with the dynamics reported in other studies on grapevine with different pot sizes and various climatic conditions (Lebon *et al*., 2006). The most obvious effect of ethanol priming was the slower drop of FTSW after priming with 250 mM for WS plants, which corresponds to a dilution of 1.4% EtOH/water (v/v) (Fig. 3B). Interestingly, a limitation of FTSW drop between two irrigation events that was also observable in WW plants pre-treated with a similar EtOH dose (Fig. 3A).

### Ethanol reduces cuttings transpiration

As pots without plants showed marginal FTSW decrease over the period of experiment due to evaporation (Supplementary Fig. S1), we assumed that all water loss mainly resulted from plant transpiration. As it was demonstrated by Bashir *et al*. (2022) in three other plant species, the EtOH priming reduced plant transpiration in grapevine (Fig. 4). The strongest effect was obtained with 250 mM EtOH. This EtOH concentration reduced transpiration in both WW and WS plants, after three days of experiment. Interestingly, the WS plants treated with this 250 mM concentration maintained a higher transpiration rate than WS controls (pre-treated with water alone) after 8 days of experiment. This effect could be beneficial in cases of combined water deficit and heat stress, as transpiration is critical to cool down leaf temperature (Costa *et al*., 2012). Even more interesting, we want to attract reader attention to the responses also obtained with the EtOH smallest dose, 0.4 mM which corresponds to 0.002% EtOH/water (v/v). Yet, the 0.4 mM treated plants displayed a slightly higher level of transpiration than controls, visible from day 8 to day 10 (Fig. 4), however it was not significant in our trial conditions. These effects induced by various EtOH doses could generate big water savings at a vineyard scale, and should definitely prompt us to test these treatments at a larger scale.

### Transpiration variations induced by ethanol follow a hormesis response

These effects obtained at low and high EtOH concentrations are logical, as EtOH is known to generate hormesis responses (Calabrese and Baldwin, 2003; Diot *et al*., 2024a). These responses harbour a bell-shaped curve as a function of increasing concentration of the priming chemical. This is quite obvious in all curves of the well-watered plants (Fig. 4). To illustrate this point, we reproduced the plots of day 10 with bell-shaped curves in Supplementary Fig. S3. The green curve highlights an increased transpiration at low ethanol doses in WW plants. Remarkedly, the water stress led to inversing the hormesis response, highlighted by the red curve in WS plants. Hormesis responses are known to be the initial parts of sigmoid curves (Kendig *et al*., 2010), and it is known that environmental conditions can modulate hormesis responses in plants (Belz and Cedergreen, 2010). These latter showed that environmental factors, such as temperature and light, modulated the stimulation of lettuce growth by low doses of a phytotoxin, parthenin. However, further experiments would be necessary to bring more explanations about modulation of grapevine transpiration in response to a combination of EtOH priming and water stress.

### Ethanol modulates the FTSWt at which transpiration decreases

The progressive soil drying led to lower affects plant transpiration from FTSWt equal to 0.44 on plants not subjected to ethanol priming (Fig. 5). This value is very close to 0.4, the value reported for grapevines in previous studies using the two-segment plateau regression procedure (Lebon *et al*., 2003, Hofmann *et al*., 2014). FTSWt values calculated using the two-segment plateau regression procedure resulting in lower values than those obtained in our study with the inverse exponential model (Kang *et al*., 2024). The response of transpiration to FTSW is employed in various dynamic crop models to monitor vineyards water status (Lebon *et al*., 2003; Valdés-Gomez *et al.,* 2009 ; Ramos *et al*., 2014; Er-Raki *et al.,* 2021), and it is worth noting that ethanol is modulating this response. Notably, priming with 250 mM EtOH resulted in an earlier regulation of transpiration with FTSWt equal to 0.67 (instead of 0.44 for 0 mM EtOH, Table 1). Ultimately, priming with 250 mM EtOH reduced transpiration and leaf growth both for WW and WS plants at first, but favoured transpiration recovery for WS plants after 8 days.

### Ethanol effects on leaf development

The hormesis effect was also observed when looking at the green leaf area at harvest in WW plants, around four weeks after trial beginning (Fig. 6) with a slight stimulation of leaf area at 2 mM, regardless of the water-deficit conditions. These results also showed that high doses of ethanol (250 mM) limited the leaf development whatever the water deficit, an effect already visible at 50 mM in WW plants. This response to exogenous ethanol has already been observed in oilseed rape by Wu *et al*. (2019). However, the water deficit induced a stronger effect than ethanol, clearly limiting leaf development in all WS cases compared to WW cases. The observation of high percentages of dry leaves, mainly in WS plants (Fig. 7), was as expected a classical plant response to water deficit (Fischer and Turner, 1978). The ethanol priming did not significantly modulate this drought effect in WS plants, in contrast with observations on cuttings with bare roots (Fig. 2). According to Hochberg et al. (2017), older basal leaves are characterised by greater connectivity in the xylem and higher vulnerability to embolism propagation, leading to the functional protection of apical younger leaves. In our case, the choice of the harvest date, based on the wilting of top leaves, which did not occur for 250 mM treatment, was probably not relevant and too late.

Nevertheless, the absence of ethanol effect on the percentage of dry leaves in WW plants (Fig. 7), always below 10%, proved that those EtOH concentrations were not toxic.

Additionally, we measured the dry weight of the root systems (Supplementary Fig. 4). Once again, we observed a hormesis effect in the WW plants with an increased root weight from 500 mg in controls to 600 mg in 2 mM EtOH treated plants, thus a +20% increase in dry weight, but this trend was not significant. Nevertheless, this boosting effect of ethanol on root development has already been observed in various plant species, such as mung bean (Bhattacharya *et al*., 1985) and oilseed rape (Wu *et al*., 2019). At higher EtOH doses, above 2 mM, the dry root weight decreased to control levels. In further experiments, ethanol should be tested in grafted plants as most grapevines are grafted nowadays.

### Potential regulations at the transcription level

In an attempt to find explanations for the differences in transpiration, we used the RNA-seq data from a recent study (Diot *et al*. 2024b), performed on Gamay cells, to screen transcript accumulation in response to an exogenous ethanol application. First, we must highlight that Gamay Freaux is genetically very close to Gamay, as it is Gamay mutant (see cultivars numbers 4377 and 4382 in https://vivc.de/). We searched particularly for markers of the metabolisms revealed by Bashir *et al*. (2022) as major causes of ethanol-induced resistance to drought: abscisic acid (ABA), aldehyde dehydrogenase (Ald DH), gamma-aminobutyric acid (GABA), neoglucogenesis, starch and sucrose (Supplementary Fig. S5). However, no marker stood out as a critical consequence of the priming by ethanol, even if a slight trend for down regulation of sucrose related genes was observed. Nevertheless, the RNA-seq data are available through the recent study (Diot *et al*., 2024b) and it may be a resource for teams looking for a particular gene or sets of genes. We also checked for regulation of calcium related gene transcripts, as calcium involvement has been described in stomatal closure (Huang *et al*., 2019), but there was no tendency for up or down regulation of calcium related genes. Diot *et al*. (2024b) observed a strong induction of small heat shock proteins (sHSPs) after ethanol priming of Gamay cells. The link between sHSPs and drought stress is known (Rizhsky *et al*., 2002; Sarkar *et al*., 2009) and Reddy *et al*. (2014) have observed the induction of class I and II sHSPs following drought stress in barley. Finally, it is worth noting that heat shock protein families, HSC70 and HSP90, are known to be involved in stomatal closure (Clément *et al*., 2011). Whether small HSPs are also involved in stomatal closure would deserve further research, but our observations: EtOH priming reduced transpiration, as shown in this article, and EtOH priming induced a strong sHSPs up-regulation, as shown in the RNA-seq data (Diot *et al*., 2024b), highlight a potential positive correlation.

## Conclusion

We showed that ethanol priming induces a slowdown of the grapevine transpiration, which was obvious with the 250 mM dose, but could also be effective at much lower dose around 0.4 mM. This latter concentration corresponds to an ethanol dilution of 0.002% in water, which would be easy and economical to implement; even though the ethanol dilution of 1.4% in water, equivalent to 250 mM, is still relatively low. Confirming the promising effects of such low doses would deserve further studies with older vines, either at greenhouse or vineyard scales. Additionally, we also observed that grapevine responds to the range of ethanol concentrations in a typical hormesis way. This clearly requires more studies to better understand this type of biochemical responses, linked to chemical priming or hormesis, which are now attracting more attention from the scientific community (Sako *et al*., 2020; Salinitro *et al*. 2021). Finally, the links between sHSPs, ethanol priming and plant resistance to drought also deserves more attention.

## Abbreviations

EtOH: Ethanol
FTSW: Fraction of Transpirable Soil Water
FTSWt: Fraction of Transpirable Soil Water threshold
HSPs: Heat Shock Proteins
NTR: Normalised Transpiration Rate
sHSPs: small Heat Shock Proteins
WS: Water Stress
WW: Well Watered

## Acknowledgments

We thank EUR TULIP and the Occitanie Region for the PhD grant to AD. We also thank M. Bouzayen and J. Pirrello for GBF team co-direction and side-funding, L. Lemonnier and D. Saint-Martin for help in cell and plant cultures.

## Author contributions

NAK, AP, PM and CC conceptualised the project and designed the research. OY provided the Gamay cuttings. NAK, BQ, PM and CC performed the cuttings experimentations. AD provided RNA-seq results obtained on Gamay cells. NAK, AD, PM and CC ran statistical tests. All co-authors contributed to writing and editing the manuscript.

## Data availability

The data underlying this article will be shared upon request to the corresponding authors.

## Supplementary Data

**Supplementary Fig. S1.**
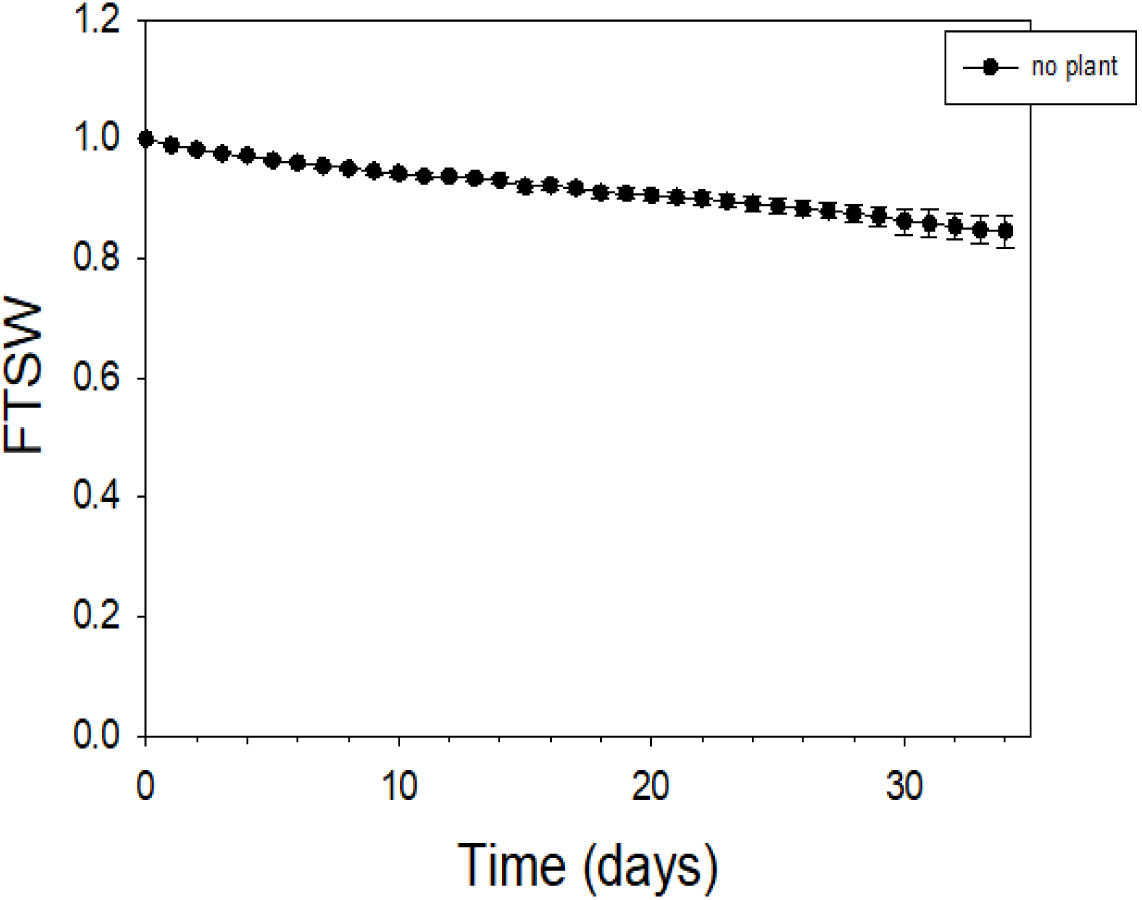
Variation of FTSW over time in pots without plants, n = 8 pots, error bars show SE.

**Supplementary Fig. S2.**
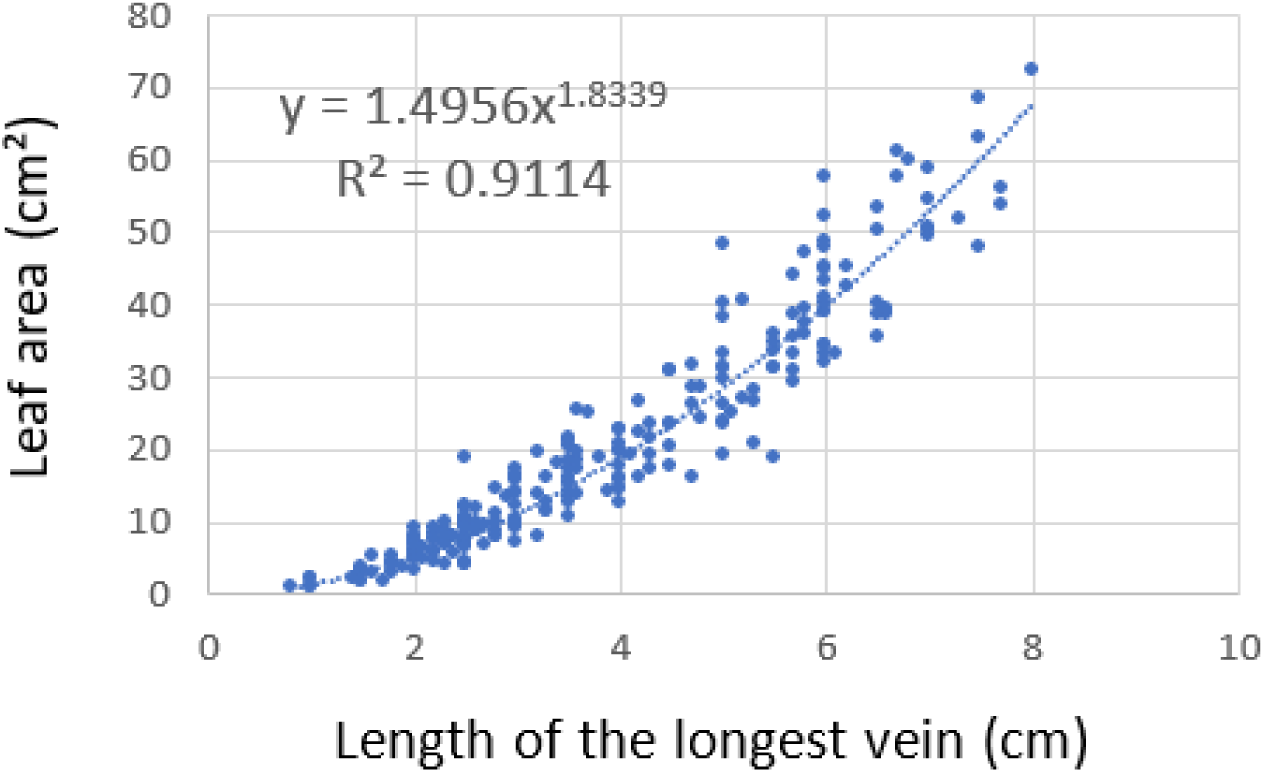
Regression between the longest vein and the leaf area, n = 233 leaves measured in 30 different Gamay one-node cuttings, dedicated to this purpose, in the same experiment and similar growing conditions.

**Supplementary Fig. S3.**
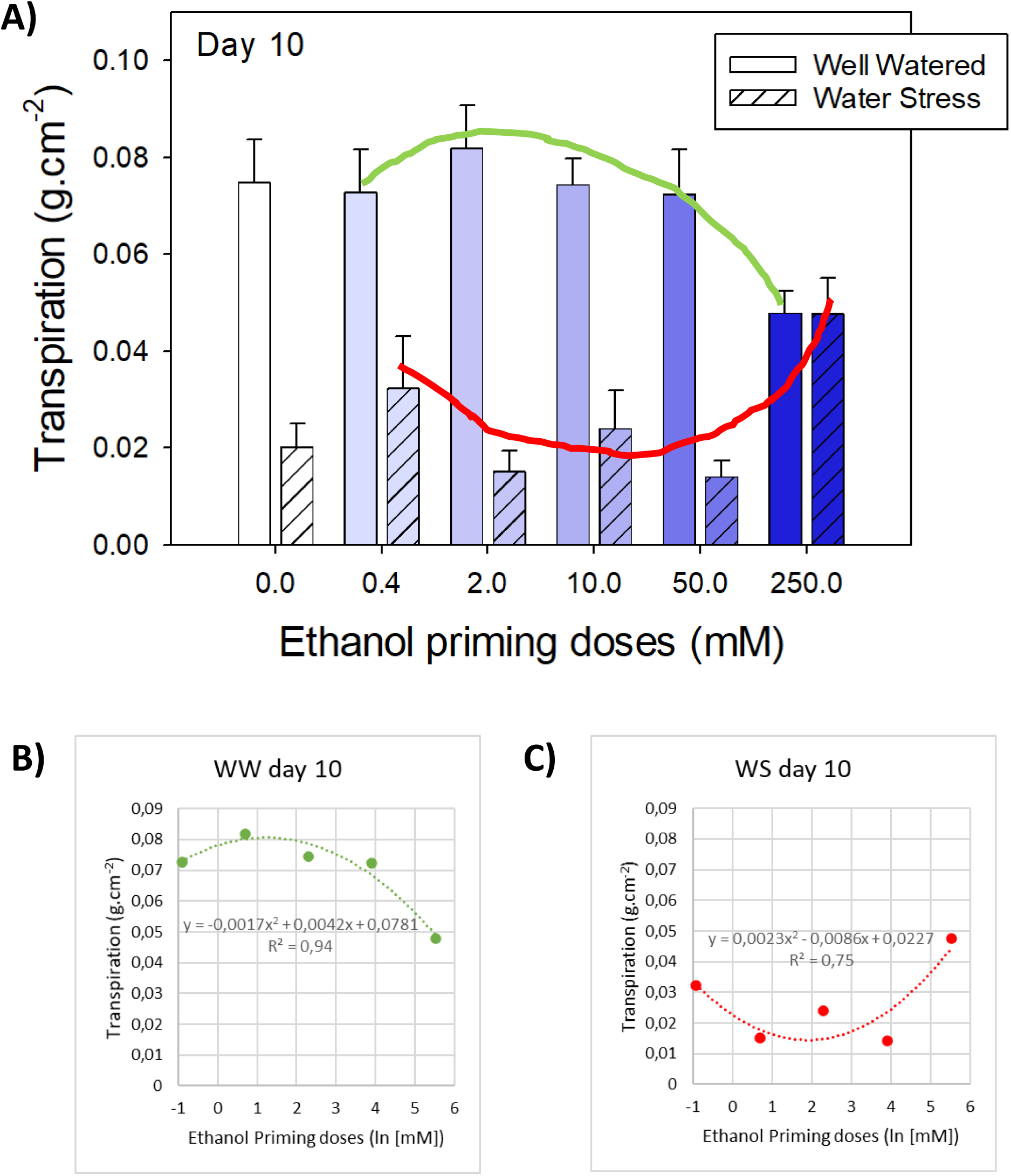
Hormesis effects on transpiration, leading to bell-shape curves, green for WW, red for WS (panel A) as a function of increasing ethanol doses. The water stress is changing the bell-shape curve, and such an environmental effect on the hormesis response has been described by Belz and Cedergreen (2010). The polynomial regressions using means (panels B and C) give an indication of the fit, to support the freehand curves.

**Supplementary Fig. S4.**
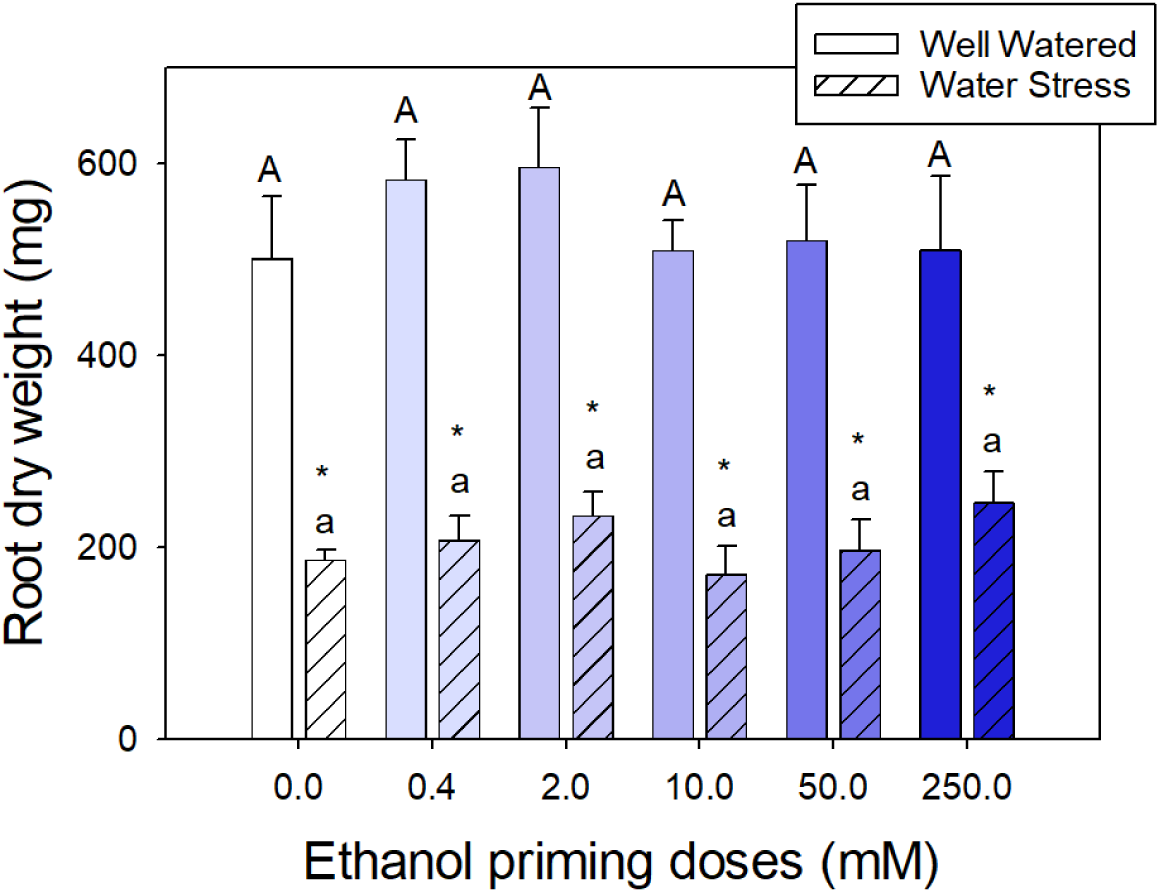
Impact of ethanol (EtOH) priming on the root development at harvest, n = 7 plants, error bars show SE. Pairwise comparisons detailed in Supp. Table S7, * shows *p* < 0.05 in comparisons of water deficit effect within each EtOH dose.

**Supplementary Fig. S5.**
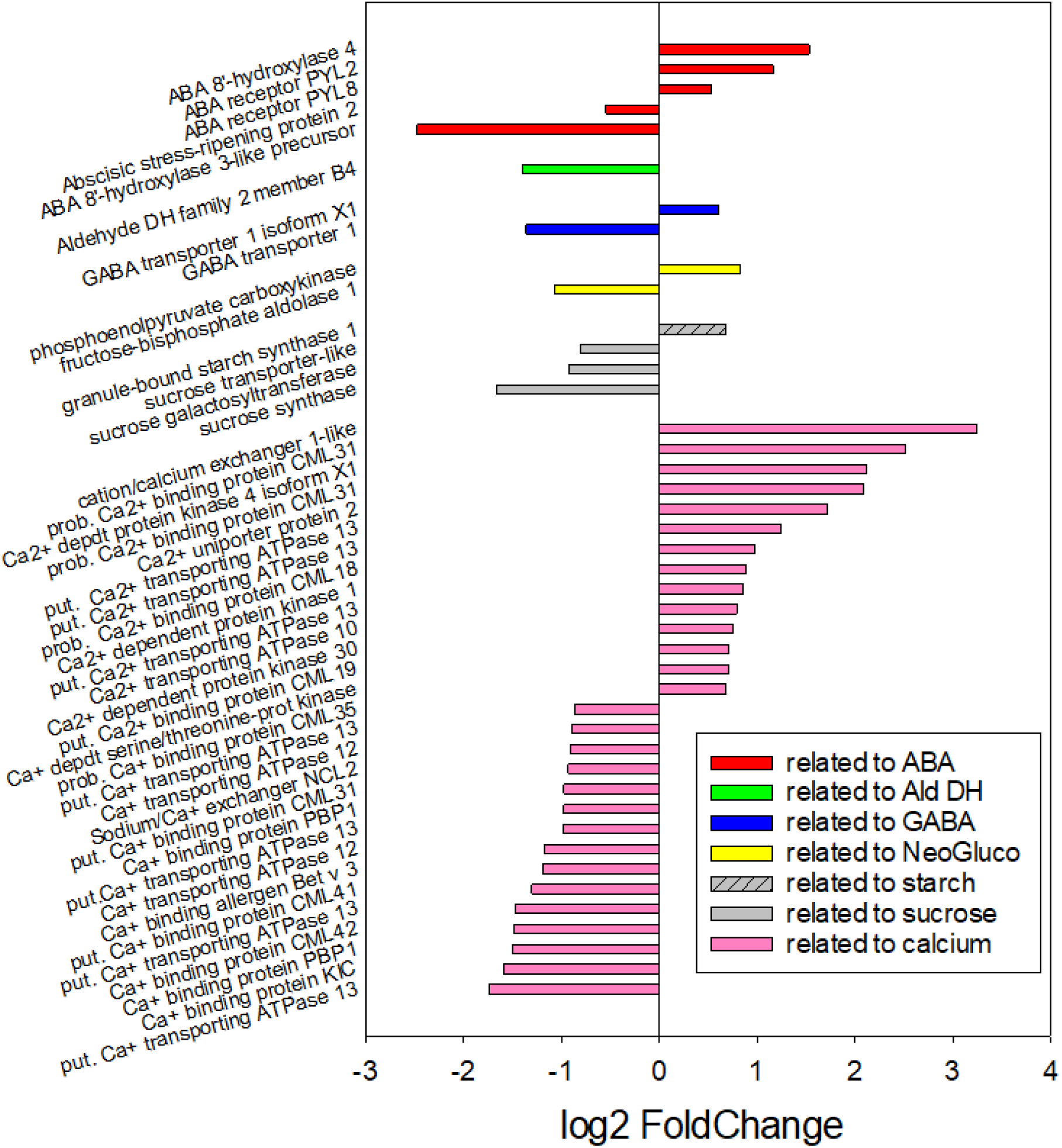
Differentially Expressed Genes linked to metabolisms related to stomatal closure, for the comparison 0 vs. 1 mM EtOH priming of Gamay cell cultures, 6 hours after treatment (see Diot *et al*., 2024b, online preprint) https://www.biorxiv.org/content/10.1101/2024.08.31.610606v2

## Supplementary Tables

**Suppl. Table S1:**
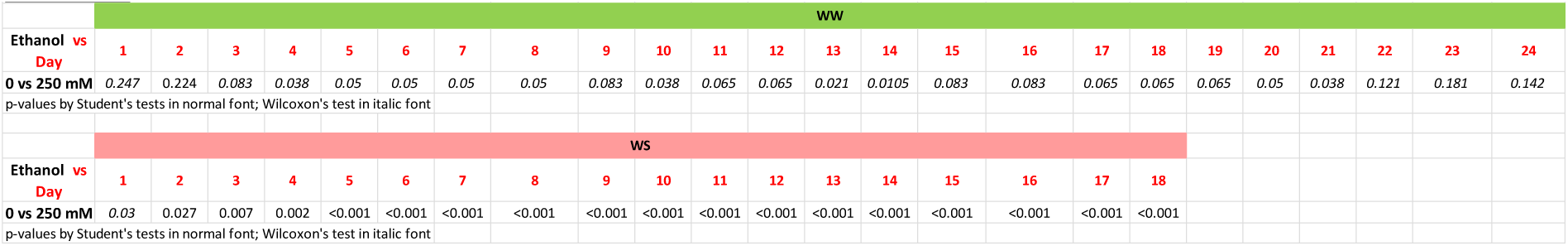
Pairwise comparisons on data of Figure 3.

**Suppl. Table S2:**
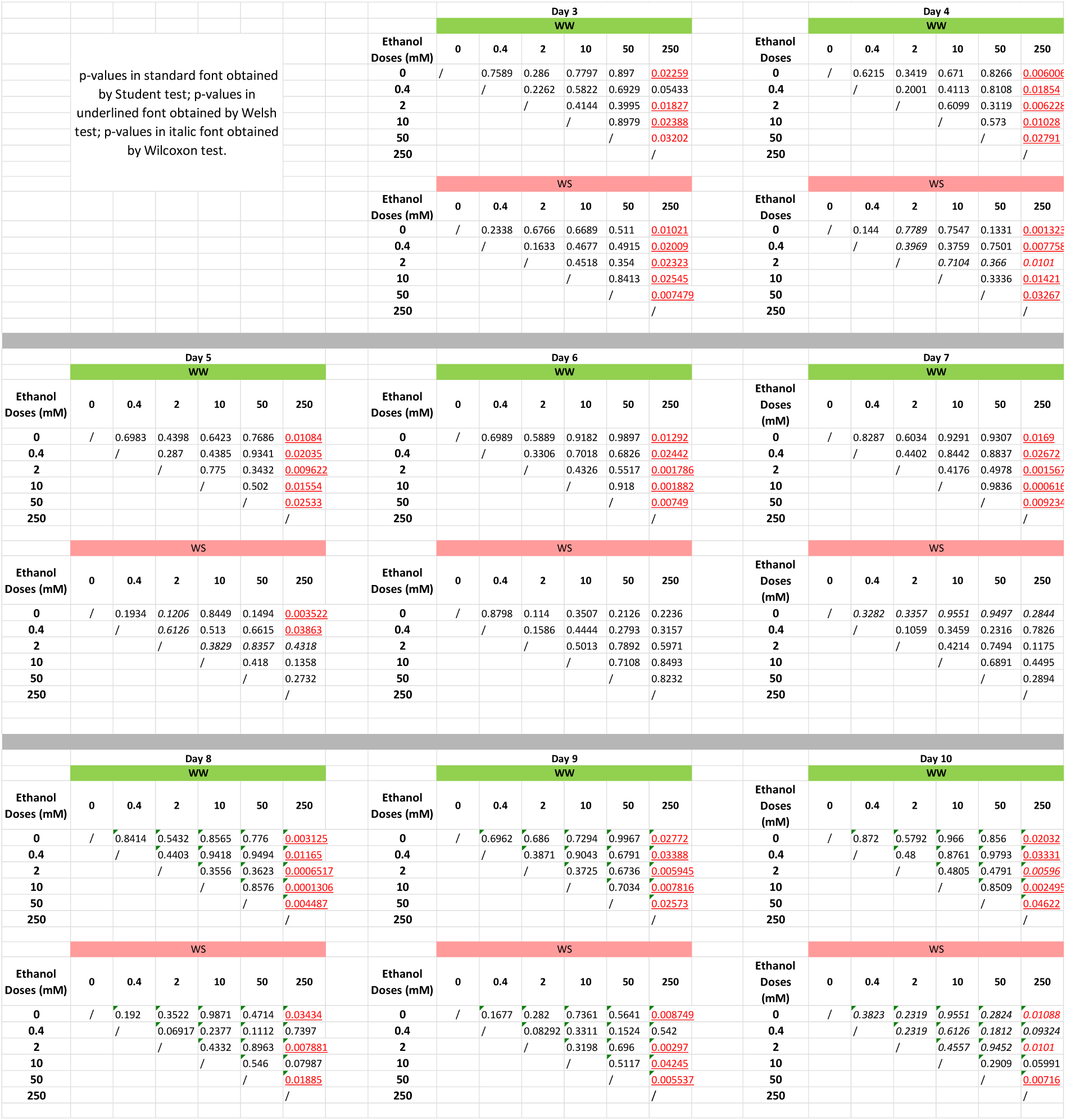
Pairwise comparisons on data of Figure 4 between EtOH doses within WW or WS.

**Suppl. Table S3:**
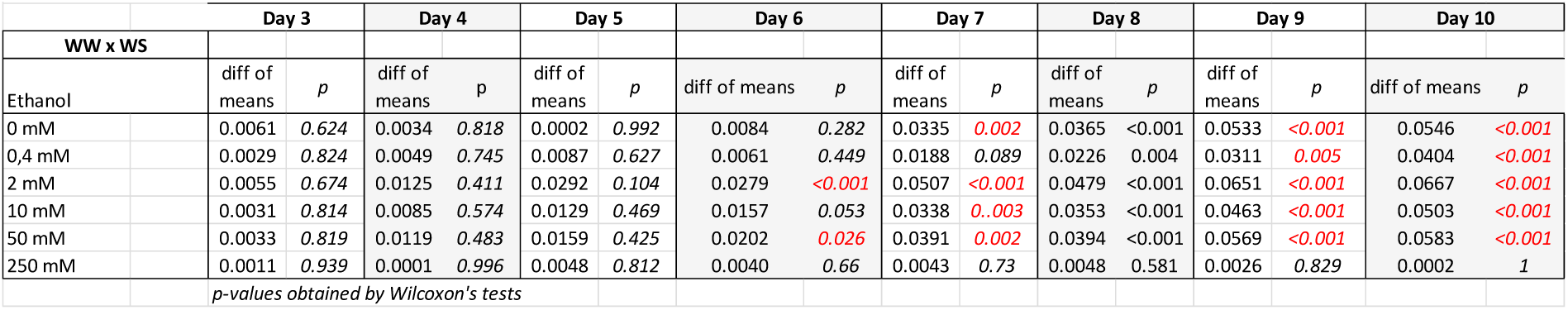
Pairwise comparisons on data of Figure 4 between WW or WS within one EtOH dose.

**Suppl. Table S4:**
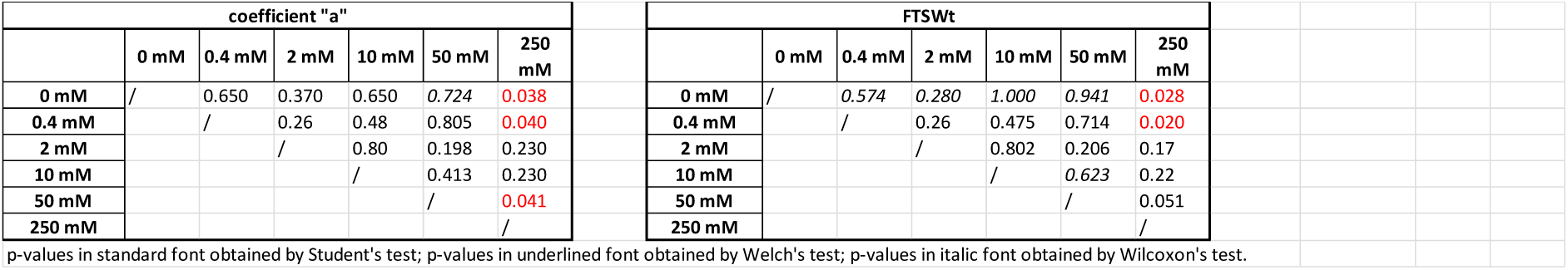
p-values of pairwise comparisons regarding “a” and FTSWt. All fitting and calculations details are given in Material and Methods.

**Suppl. Table S5:** Pairwise comparisons on data of Figure 6 between EtOH doses within WW or WS.

| WW |  |  |  |  |  |  | between WW vs WS, within EtOH |  |
| --- | --- | --- | --- | --- | --- | --- | --- | --- |
| Ethanol Doses (mM) | 0 | 0.4 | 2 | 10 | 50 | 250 |  |  |
| 0 | / | 0.9977 | 0.4614 | 0.7649 | 0.103 | 0.03075 | WW x WS |  |
| 0.4 |  | / | 0.4592 | 0.7666 | 0.07529 | 0.03882 |  |  |
| 2 |  |  | / | 0.2555 | 0.0067 | 0.0025 | Ethanol | p |
| 10 |  |  |  | / | 0.1085 | 0.02954 | 0 mM | 0.026 |
| 50 |  |  |  |  | / | 0.3191 | 0.4 mM | 0.003 |
| 250 |  |  |  |  |  | / | 2 mM | 0.002 |
|  |  |  |  |  |  |  | 10 mM | 0.014 |
|  |  |  |  |  |  |  | 50 mM | 0.011 |
|  |  |  |  |  |  |  | 250 mM | 0.005 |
| WS |  |  |  |  |  |  |  |  |
| Ethanol Doses (mM) | 0 | 0.4 | 2 | 10 | 50 | 250 |  |  |
| 0 | / | 0.4445 | 0.9251 | 0.9289 | 0.3894 | 0.2397 |  |  |
| 0.4 |  | / | 0.3054 | 0.4552 | 0.9435 | 0.3469 |  |  |
| 2 |  |  | / | 0.8337 | 0.2577 | 0.1095 |  |  |
| 10 |  |  |  | / | 0.3988 | 0.2486 |  |  |
| 50 |  |  |  |  | / | 0.2572 |  |  |
| 250 |  |  |  |  |  | / |  |  |

p-values in standard font obtained by Student's test; p-values in underlined font obtained by Welch's test; p-values in italic font obtained by Wilcoxon's test.

**Suppl. Table S6:** Pairwise comparisons on data of Figure 7 between EtOH doses within WW or WS.

| WW |  |  |  |  |  |  | between WW vs WS, within EtOH |  |
| --- | --- | --- | --- | --- | --- | --- | --- | --- |
| Ethanol Doses (mM) | 0 | 0.4 | 2 | 10 | 50 | 250 |  |  |
| 0 | / | 0.9351 | 0.4984 | 0.7403 | 0.8713 | 0.9351 | WW x WS |  |
| 0.4 |  | / | 0.599 | 0.6282 | 0.9233 | 0.99 |  |  |
| 2 |  |  | / | 0.2952 | 0.8255 | 0.599 | Ethanol | p |
| 10 |  |  |  | / | 0.6291 | 0.6282 | 0 mM | < 0.001 |
| 50 |  |  |  |  | / | 0.9233 | 0.4 mM | < 0.001 |
| 250 |  |  |  |  |  | / | 2 mM | < 0.001 |
|  |  |  |  |  |  |  | 10 mM | < 0.001 |
|  |  |  |  |  |  |  | 50 mM | < 0.001 |
|  |  |  |  |  |  |  | 250 mM | 0.002 |
| WS |  |  |  |  |  |  |  |  |
| Ethanol Doses (mM) | 0 | 0.4 | 2 | 10 | 50 | 250 |  |  |
| 0 | / | 0.6421 | 0.7548 | 0.8899 | 0.7548 | 0.09923 |  |  |
| 0.4 |  | / | 0.8701 | 0.5326 | 0.8701 | 0.1133 |  |  |
| 2 |  |  | / | 0.6409 | 0.99 | 0.0713 |  |  |
| 10 |  |  |  | / | 0.6409 | 0.0527 |  |  |
| 50 |  |  |  |  | / | 0.0713 |  |  |
| 250 |  |  |  |  |  | / |  |  |

p-values in standard font obtained by Student's test; p-values in underlined font obtained by Welch's test; p-values in italic font obtained by Wilcoxon's test.

**Suppl. Table S7:**
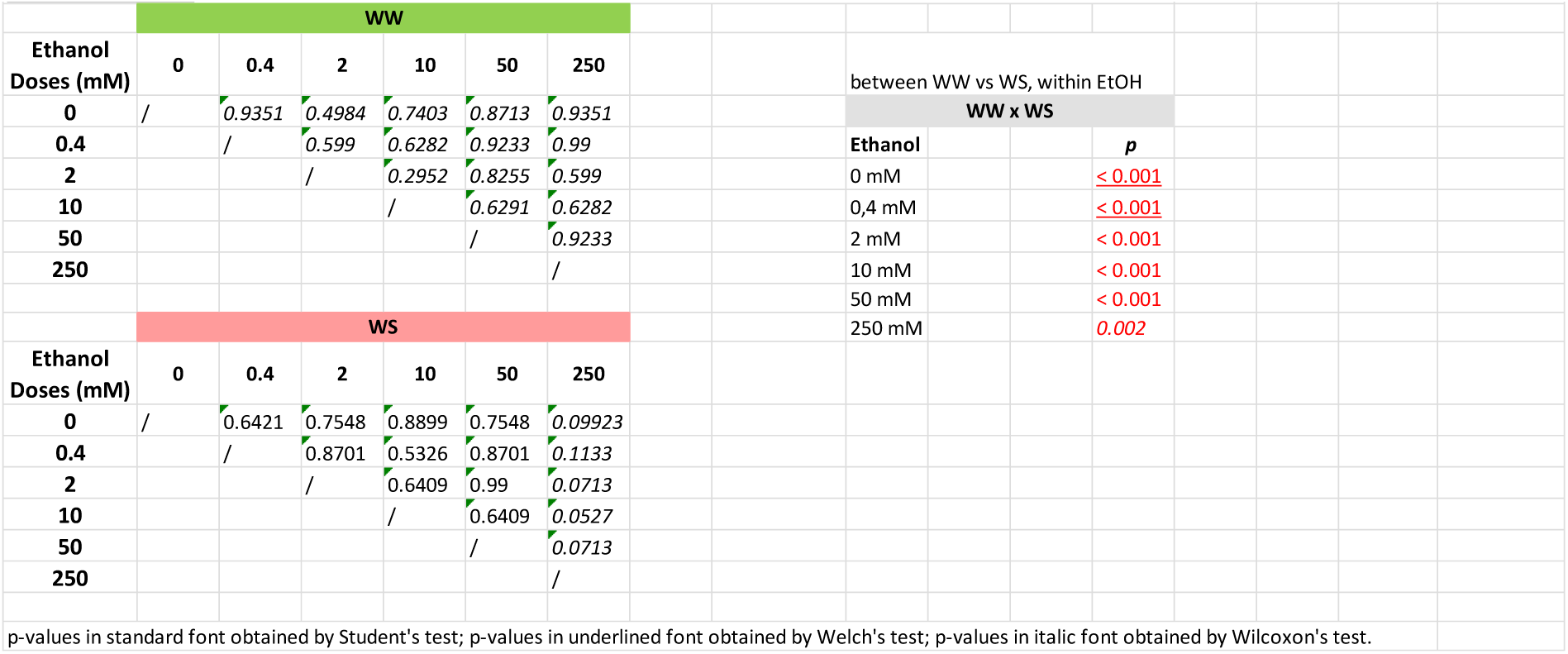
Pairwise comparisons on data of Supp. Fig. S4 between EtOH doses within WW or WS.

